# The centrality of ecotones: How scale, sex, and ontogeny shape the spatial ecology of a solitary carnivore

**DOI:** 10.64898/2026.05.01.722308

**Authors:** Paul Glover-Kapfer, Qian Song, John Erb

## Abstract

**Context:** Animals balance resource acquisition with risk mitigation. These trade-offs are rarely uniform, being mediated by spatial scale, demographic traits, and environmental constraints. Understanding these divergent spatial behaviors is critical for management across human-dominated landscapes.

**Objectives:** We investigated how sexual dimorphism and ontogeny interact with landscape structure to influence scale-dependent resource selection. Specifically, we sought to determine how these demographic factors mediate spatial trade-offs between optimal foraging habitats, top-down intraguild predation risk, and bottom-up severe winter weather.

**Methods:** We examined the spatial ecology of a solitary carnivore, the bobcat (*Lynx rufus*), across a heterogeneous, human-modified landscape in northern Minnesota, USA. Using spatial data derived from harvested adult and juvenile individuals, we evaluated multi-scale selection relative to land cover, structural ecotones, intraguild predator activity, and winter severity.

**Results:** Habitat selection was scale-dependent and partitioned demographically. Whereas bobcats universally selected for ecotones and avoided homogeneous open habitats at fine scales, responses to other features diverged by sex and age. Females actively avoided areas with high coyote activity and freezing temperatures; males exhibited high risk tolerance, apparently indifferent to coyote activity and tolerant of freezing temperatures. We identified a distinct ontogenetic spatial shift among females. Subordinate juveniles were competitively excluded from optimal natural ecotones, forcing them into riskier, anthropogenic agricultural edges. In contrast, adult females optimized foraging opportunities by selecting productive ecotones at the intersection of woody vegetation and semi-natural grasslands.

**Conclusions:** Our findings demonstrate that habitat selection is not a static species-level trait, but instead a dynamic process resulting from the interaction between ontogeny, sex, and landscape heterogeneity. The reliance of vulnerable demographic groups on marginal or anthropogenic habitats highlights how human land-use changes can inadvertently produce ecological winners and losers within the same species. Consequently, landscape management and conservation planning for solitary carnivores must shift from broad, population-wide habitat prescriptions to strategies that explicitly accommodate the divergent spatial requirements of specific demographic cohorts.

## Background

To navigate heterogeneous and dynamic landscapes, animals must balance resource acquisition with risk management, and this trade-off defines animal space use. Because resources and risks vary across a hierarchy of spatio-temporal scales, so too do patterns of space use (Johnson 1980) as animals seek to exploit resources while for many also avoiding becoming one themselves. Animals may respond to persistent or predictable factors at coarse scales, such as regional climate gradients or extensive human land use, and exhibit more fine-scale responses to localised and less predictable resources and threats, such as the dynamic distribution of prey or predators (Jensen et al. 2024). For example, species distributional patterns are often related to climatic patterns, whereas localised selection of micro-habitats may be determined by weather. Understanding the hierarchy of space use is therefore fundamental to understanding species ecology and the development of mechanistic understandings of their response to environmental change (Gomez et al. 2025).

The trade-off between resource acquisition and risk mitigation is further complicated by intraspecific physiological and physical variation, which dictates not only energetic requirements but also determines resource availability and vulnerability. Sexual dimorphism is perhaps the most visible manifestation of this variation.

Many vertebrate species exhibit pronounced sex-based dimorphism in body size, physiology, and behavior, resulting in divergent energetic demands (Taylor et al. 1972), sensitivities to risk, ecological roles (Gissi et al. 2024), and resource use (Andrén and Liberg 2015; Adams et al. 2017). Although the relative role of natural versus sexual selection in shaping sexual dimorphism is contextual, these differences frequently result in sex-specific selective pressures and life history strategies. As such, assuming a single, population-wide habitat-selection pattern obscures important heterogeneity and can lead to incorrect inferences about the environmental drivers of space use (Conde et al. 2010).

Sexual dimorphism can be particularly pronounced among the Carnivora, with body masses differing by up to 320% (Tombak et al. 2024), the consequences of which are multiple. For example, males may face a decreased risk of predation (Wengert et al. 2014), interspecific competition (Bisazza et al. 1996), and intraspecific aggression (Oddie 2000; Serrano-Meneses et al. 2007) because of their larger size, and therefore more readily access resources. Conversely, larger body size imposes greater energetic costs for growth and maintenance, which can necessitate reliance on high-value, potentially dispersed resource patches. This requires more extensive movements or resource defense – behaviors that may increase exposure to risk via intra- or interspecific competition or predation (Norrdahl and Korpimäki 1998; Thurfjell et al. 2017). Synergy between larger body size and extensive movements may also increase exposure to human-caused mortality (e.g., roadkill, hunting), especially for species targeted as trophies (Ciuti et al. 2012; Rivrud et al. 2013; Allen et al. 2018; Bischoff et al. 2018).

As solitary, obligate carnivores, members of the family Felidae exhibit pronounced sexual size dimorphism (Law 2019), which translates into a substantial degree of ecological dimorphism. Empirical examinations of sex-specific diets support this supposition, demonstrating that intersexual trophic niche partitioning in felids is substantial (Litvaitis et al. 1986; McLean et al. 2005; Meiri et al. 2005). Beyond dietary partitioning, this divergence extends directly into spatial ecology and producing sex-specific differences in habitat selection (Bauer et al. 2025).

A largely missing yet critical component of the mechanisms responsible for observed variation in space use is the ontogenetic shift in space use that occurs as individuals grow and mature. Many individuals in early stages of development are subject to increased risk from not only predation, but also interspecific and intraspecific competition. Consequently, their space use should theoretically emphasise risk-avoidance strategies. As they mature physically and gain experience, risk from these sources declines, resulting in an ontogenetic shift toward an emphasis on optimal foraging strategies. This shift may be particularly pronounced for species for which maturation is associated with the transition from a transient or dispersal phase to residency, as each phase imposes distinct ecological costs and benefits. Because transients lack detailed familiarity with their environs, they may experience elevated risk (Yoder et al. 2004), thereby favouring behaviors that emphasise safety.

Residents, however, benefit from familiarity with their local environment, thereby reducing risk, and should pivot to behaviors that emphasise territorial establishment and defense and energy acquisition.

The bobcat (*Lynx rufus*) provides an ideal model to investigate the role of ontogeny and sex in determining space use because it is affected by both top-down and bottom-up pressures. As a mesocarnivore, bobcats experience intraguild competition and potential predation from larger sympatric carnivores (Gipson and Kamler 2002; Hass 2009), and because of their relatively high foot-loading (mass-to-footpad area ratio; Buskirk et al. 2000) and lower critical temperature (Mautz and Pekins 1989) are susceptible to winter severity in the form of deep snow and freezing temperatures. How individual bobcats navigate these competing risks and rewards across different spatial scales remains poorly understood, particularly with regard to the modifying effects of sex and age.

To address these questions, we evaluated how sex and female ontogeny influence space use in bobcats in Minnesota across three spatial scales (landscape, home range, and local). We assessed support for three hypotheses predicting how landscape heterogeneity mediates risk-resource trade-offs: (1) Scale-dependent ecotone selection: Across all scales, all individuals will select structurally complex ecotones for ambush foraging while avoiding homogeneous, open agricultural matrices. (2) Sex-specific landscape permeability: At local scales, reproductive females will restrict space use to mitigate risk from intraguild competition, while avoiding severe weather at broader scales. Conversely, larger males will exhibit greater tolerance for climatic and competitive risks to maximize access to reproductive opportunities. (3) Ontogenetic spatial exclusion: Female space use will shift with age. Immature juveniles will be competitively excluded from optimal natural ecotones, forcing them into, anthropogenic edges, whereas adult females will dominate highly productive semi-natural ecotones.

## Methods

### Data collection and study area

We worked with the Minnesota Department of Natural Resources (MNDNR) to collect location data on bobcats harvested during the 2008-2010 winters. Hunters and trappers that harvest bobcats in Minnesota are required to register with the MNDNR within 48 hours of the close of the season, at which time the DNR collects data on sex, date, township, and method of harvest. Additional data requested for this study included the section where the bobcat was harvested. Harvested bobcats were sexed via visual examination of external and if necessary, internal reproductive tracts. Female bobcats were aged using cementum annuli counts, and males were classed as young of the year (∼6 months of age) or adult based on the presence/absence of an open root foramen (Crowe 1975). Given that bobcat parturition primarily peaks between May and June (Crowe 1975) and the harvest season spanned from late November to early January, we estimated the biological age of each individual at the time of harvest as the cementum annuli count plus approximately six months.

We acknowledge that using harvest locations may introduce bias, as these data represent the intersection of bobcat occurrence and the spatial distribution of harvest effort. To explicitly account for this spatial bias, landscape covariates associated with human access (e.g., road densities) were evaluated in our candidate model sets. Rather than categorically excluding these covariates, we critically evaluated the directionality of their relationships to decouple true biological habitat selection from anthropogenic sampling bias. For example, a positive association between bobcat use and dirt road density would be equivocal, as it could reflect either true habitat selection or concentrated hunter effort. Conversely, a negative relationship provides a conservative, unambiguous signal of behavioral avoidance, demonstrating that bobcats are actively selecting against these features despite the presumed higher probability of human detection and access. Furthermore, we argue that it is important to contextualize this limitation within the broader scope of carnivore research. Most non-invasive sampling frameworks, including camera trapping for density or occupancy estimation, as well as track surveys, are frequently deployed along established linear features or access routes to maximize detection and survey convenience, and therefore similarly deviate from the ideal randomized sampling. Moreover, while modern GPS telemetry frequently yields high-volume datasets, those locations are heavily autocorrelated and typically represent the behaviors of a small number of individuals. By utilizing harvest locations, our dataset captures over 600 spatially distinct, independent individuals, providing a robust, population-level sample size that avoids spatial pseudoreplication.

Bobcat range in Minnesota is primarily limited to the northern, forested half of the state. We delineated the study area with a 100% minimum convex polygon (MCP; Mohr 1947) encompassing all bobcat locations (Fig. 1) in northern Minnesota.

**Fig. 1.**
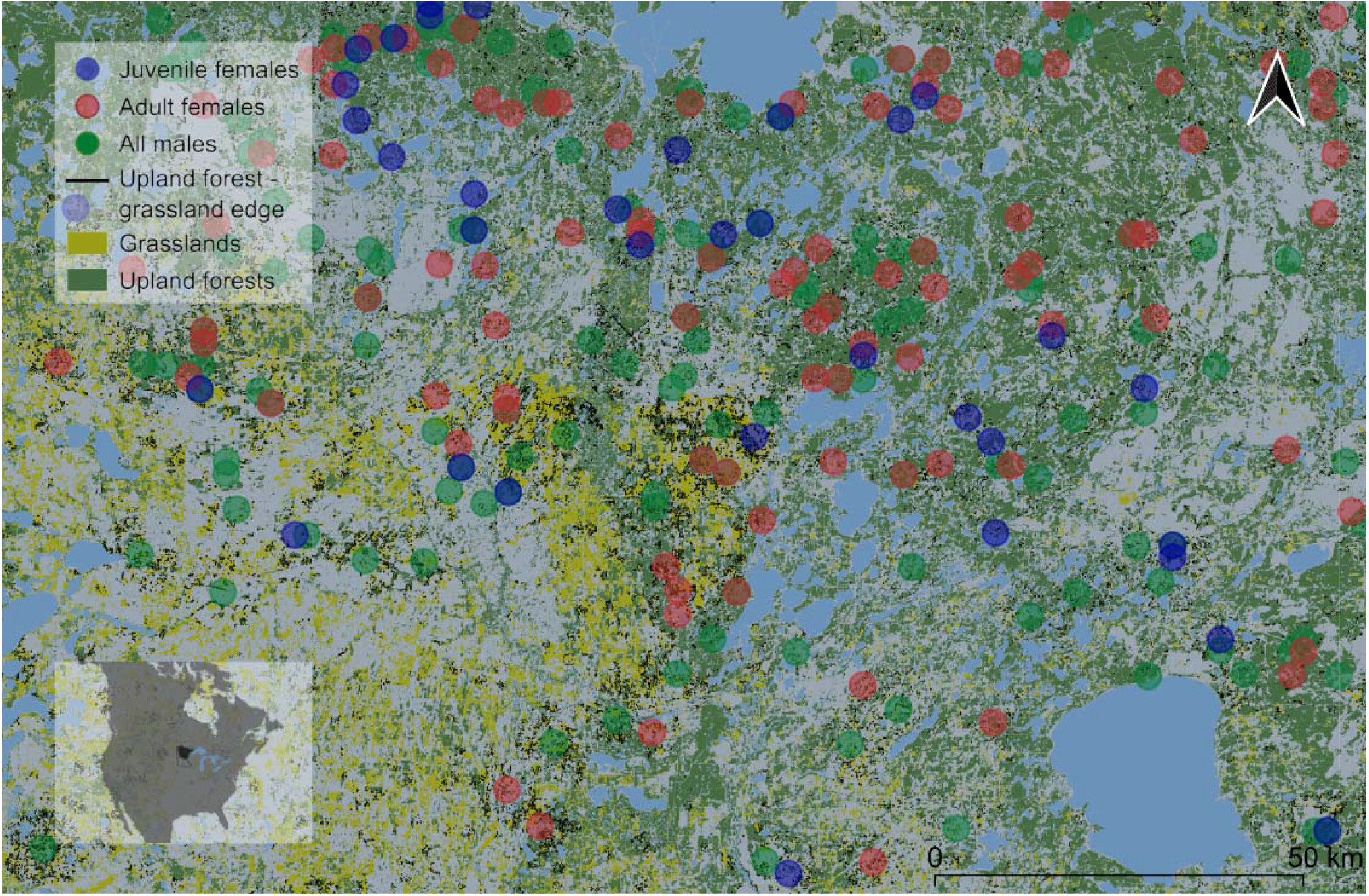
Zoomed in portion of the study area in northern Minnesota, demonstrating bobcat locations relative to the upland forest-grassland edge ecotone. Points indicate individual locations categorised by demographic groups: all males, juvenile females, and adult females. The bottom-left inset indicates the location of the study area within North America.

The MCP encompassed approximately 116,859 km2, the majority of which occurred within the Laurentian mixed forest province (Cleland et al. 1997). Dominant land cover in the study area consisted primarily of conifer (*Picea* spp., *Pinus* spp., *Thuja occidentalis*) and deciduous (*Betula papyrifera, Populus* spp., *Fraxinus* spp.) forests, bogs and swamps (71%) intermixed with agricultural fields (6%), pasture (7%), open water (7%) and rural, suburban and urban development (3%). Topography was characterized by gently rolling hills. Elevation generally increases from the southwest to the northeast and ranges from 180 to 590 m above sea level. Climate in the study area was continental with cold dry winters and hot humid summers. Normal mean temperatures range from -15° to -13° C during the winter (December to February) and 17° to 20° C during the summer (June to August). Normal mean precipitation ranges from 15 to 30 cm during the winter and 25 to 33 cm during the summer.

### Data and variable preparation

We prepared an exhaustive set of explanatory variables based on the literature and knowledge of bobcat ecology to assess sex- and ontogenetic-based differences in habitat use across spatial scales. We were particularly interested in top-down and bottom-up predictors owing to their value to understanding the mechanisms underlying bobcat spatial ecology and their transferability to other regions and species.

#### Climate, prey, and intraguild competition variable derivation

Bobcats in Minnesota are near the northern edge of their distribution and evidence suggests the northern edge of their range is defined by winter severity (citation). To assess the potential of winter weather conditions to effect bobcat spatial ecology, we included variables representing the proportion of days when the maximum temperature failed to exceed the bobcats lower critical temperature (-2.2°C; Mautz and Pekins 1989), and the proportion of days when snow depth was > 15 cm, a threshold based on observations of bobcat avoidance of deep snow (McCord 1974) and their high foot-loading (Buskirk et al. 2000) for each year of the study. These thresholds hypothetically represent the limits at which thermoregulatory costs and energy expenditure significantly increase for bobcats.

We generated winter weather conditions (name them probably) using daily weather records from 87 weather stations (MNDNR 2026) distributed throughout bobcat range, and random forest regression because of its proficiency at modelling complex, non-linear relationships. We modelled weather station data as a function of year, elevation, geographic coordinates, and proportion of a 2.5, 5.0, 7.5, and 10.0 km buffers around each weather station consisting of upland forest, development, or open water (Minnesota Land Cover Classification System values 105-107, 1-100, and 103 respectively; MNDNR 2014) for the months of November and December, corresponding approximately with our sampling dates, and excluded collinear variables from inhabiting the same model (|*r*| > 0.7). We evaluated model performance using Out-of-Bag *R*^2^ (hereafter *R*^2^) and Mean Squared Error, with *R*^2^ values for relevant predictors ranging from 0.76 to 0.83 (Table S1). These values were then projected onto the entirety of bobcat range at the same scale as bobcat locations.

We took a similar approach to modelling the relative activity of coyote and snowshoe hare. The MNDNR conducts annual standardised track surveys following fresh snowfall along set forest roads and records the presence or absence of species within each 0.8 km segment (Erb 2019a). We buffered these routes by 2.5, 5.0. 7.5, and 10.0 km and compared the proportion of segments along a survey route with coyote or snow hare to a suite of explanatory variables contained within each buffer, including proportion land cover type, land cover patch isolation, patch density, and mean patch size (using the landscapemetrics package; Hesselbarth et al.

2019), ecotone density, snow depth (as estimated above), and the density of dirt and gravel roads. We used random forest regression with 500 trees to explore all possible variable combinations, with a maximum *k* = 5. We chose models based on the highest *R*^2^ and whether the resulting model was biologically realistic. Because coyote tracks were too rare in the data to develop year-specific results, we combined all years for the coyote model. Results indicated that the most predictive variables for hare varied across years, and resulting models for both explained a reasonable amount of variation (snowshoe hare mean *R*^2^ = 0.56; coyote *R*^2^ = 0.43; Table S2).

#### Variable selection and model development

Prior to RSF development, we evaluated multicollinearity among environmental and biotic predictors using a Pearson correlation matrix, and precluded pairwise variables with |r| ≥ 0.7 from inhabiting the same model. We further refined the candidate set by removing variables that exploratory analyses indicated shared a biologically implausible relationship with bobcat space use or that was collinear with another variable that was better supported by the literature. This resulted in a reduced suite of 20–25 predictors per model for each scale (Tables S3-S5).

To test our hypotheses, we partitioned the dataset into distinct demographic cohorts based on age and sex. To establish a population-level baseline of general spatial ecology, we first modelled habitat selection using all bobcats (≥ 1-year-old) regardless of sex. We then stratified these data by sex to evaluate intersexual divergence. Finally, to investigate ontogenetic shifts, we subdivided the female cohort into juveniles (1-year-olds) and adults (≥ 2-years-old). Given the May–June parturition peak and the late-November-early-January harvest season, these classes correspond to biological ages of approximately 18 months and ≥ 30 months, respectively. We selected these thresholds because they closely align with major life-history transitions, specifically dispersal (Kamler et al. 2000; Janecka et al. 2007; Hughes et al. 2019) and the onset of reproduction (Crowe 1975). However, because harvested males were broadly classified only as either dependent kittens or adults, we were unable to investigate ontogenetic shifts in male space use. Consequently, direct comparisons between specific female age classes and the pooled male cohort should be interpreted with caution.

We assessed bobcat habitat selection across three scales, corresponding to second- and third-order selection as defined by Johnson (1980). For our landscape model we compared used locations to approximately ten-fold available locations of equal area, randomly distributed throughout a 100% MCP encompassing all recorded locations, excluding open water. This model characterises the placement of home ranges relative to the broader bobcat range in Minnesota. At this scale, we used Generalized Linear Models (GLMs) with a binomial distribution and a logit link function to model selection, with coefficients representing the change in the log-odds of bobcat use per unit increase in standardised predictors. For our home range model, we defined availability by buffering each used location to an area equivalent to the mean home range size for bobcats in the Great Lakes region (Male: 59.1 km^2^; Female: 28.8 km^2^; Fuller et al. 1985; Lovallo and Anderson 1996; Kapfer 2014) resulting in a one-to-one used-to-available comparison. For the local model, we compared used locations to approximately 10 random available locations of equal area within these buffered boundaries. To account for the hierarchical structure and varying availability, we employed stratified Conditional Logistic Regression (clogit) from the survival package (Therneau 2026) in a case-control framework.

#### Model selection and assessment

For each scale, we conducted an exhaustive search of all possible additive combinations of 1–5 predictors. We identified a competitive model set using a threshold of ΔAICc < 2 (Burnham and Anderson 2002), after which we applied model averaging to calculate weighted parameter estimates across the top-performing models. Variable significance was primarily assessed via Wald statistics, but in instances where quasi-complete separation led to inflated standard errors, we utilised Likelihood Ratio Tests to confirm variable importance.

We evaluated model performance using 5-fold stratified cross-validation with an 80:20 training-to-test data split. We assessed predictive accuracy using test and training Area Under the Receiver Operating Characteristic Curve (AUC), or the conditional logistic regression equivalent, test and training Concordance. Both metrics provide an equivalent measure of a model”s ability to discriminate between used and available locations (Harrell et al. 1996), with values > 0.7 generally indicating good to strong predictive performance (Hosmer et al. 2013). All analyses were conducted in R (version 4.5.1; R Core Team 2021).

## Results

From 2008 to 2011, we collected a total of 320 locations for males, and 285 locations for females > one year of age. The total area encompassed by all the locations was approximately 110,954 km2. This represents approximately 85% of the total range of bobcats in Minnesota as determined by a 100% MCP encompassing all harvest records from 2001-2023 (130,423 km^2^). Mean area of sections where bobcats were harvested was 2.54 km^2^ (SE = 0.06).

All pairwise correlations were below our threshold of |*r*| ≤ 0.7, indicating no significant redundancy among variables, although we removed upland forest riparian corridors from our original slate of predictors because inclusion resulted in complete statistical separation and subsequent investigation indicated that while this habitat type was present at 49.3% of available locations, it was completely absent at juvenile and adult female locations.

### Model Performance and Validation Results

Across all three spatial scales, our models exhibited high predictive accuracy and discriminatory power, and the 5-fold stratified cross-validation confirmed the robustness of these results, with minimal mean differences between training and test AUC values (Δ < 0.05, Table S3), suggesting models were parsimonious while retaining predictive performance. Our landscape-scale models had test Concordance values ranging from 0.79 to 0.83, indicating a high degree of success at predicting the broad placement of bobcat home ranges across northern Minnesota. At the local level, test Concordance values for our models comparing used to available locations within a home range buffer ranged from 0.70 to 0.75. Models at the home range scale produced our highest predictive accuracy, with test AUC values ranging from 0.87 to 0.91.

Model convergence was consistent across all spatial scales and demographic groupings. We observed a low degree of model uncertainty, with the number of competitive top models (ΔAICc < 2) remaining low and relatively uniform across all scenarios (x□ = 1.73, range: 1–3), indicating that our top-ranked models adequately described resource selection.

### Similarities and differences in habitat selection

Broadly, our resource selection functions revealed that bobcat spatial ecology is highly scale-dependent and partitioned across demographic cohorts (Fig. 2).

**Fig. 2.**
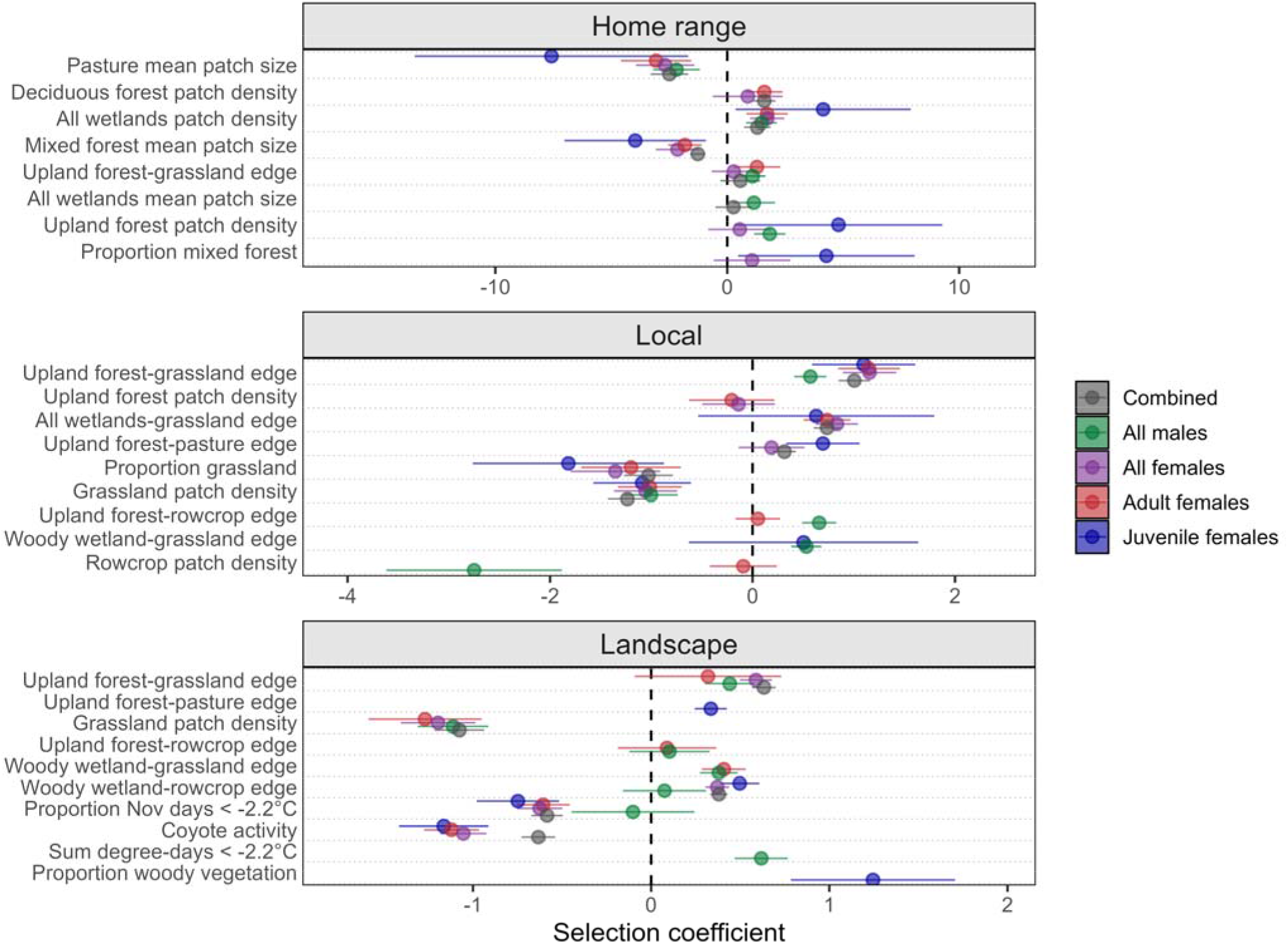
Standardised coefficient estimates (points) and 95% confidence intervals (horizontal lines) for predictors of bobcat resource selection across three hierarchical spatial scales: home range, local, and landscape. The vertical dashed line at zero represents a neutral effect, or a lack of selection, with estimates > 0 indicating positive selection and estimates < 0 indicating avoidance.

Of the 27 covariates retained in the competitive models across the 3 analyses, only four were important for all demographic groups. Pasture mean patch size at the home range scale and grassland patch density at the local scale were universally avoided, and the patch density of all wetlands at the home range scale and upland-forest grassland edge at the local scale were universally selected (Fig. 3). Outside of these shared baselines however, the influence of top-down predation risk, bottom-up thermal constraints, and anthropogenic land use diverged sharply by sex and ontogeny at coarser spatial scales. For example, although coyote activity was included in the global model of all bobcats, it was not included in any of the competitive models for males, whereas it was negatively associated with adult and juvenile females (Fig. 4). Similarly, while adult and juvenile females showed a different sensitivity to freezing temperatures, males showed the opposite response with the probability of use increasing as temperatures got colder (*β* = 0.62, *p* < 0.001 for males vs. *β* = -0.63, *p* < 0.001 for all females combined). Remaining parameter estimates, standard errors, and 95% confidence intervals for all competitive models are detailed in the Supplementary Information (Tables S7–S9, respectively).

**Fig. 3.**
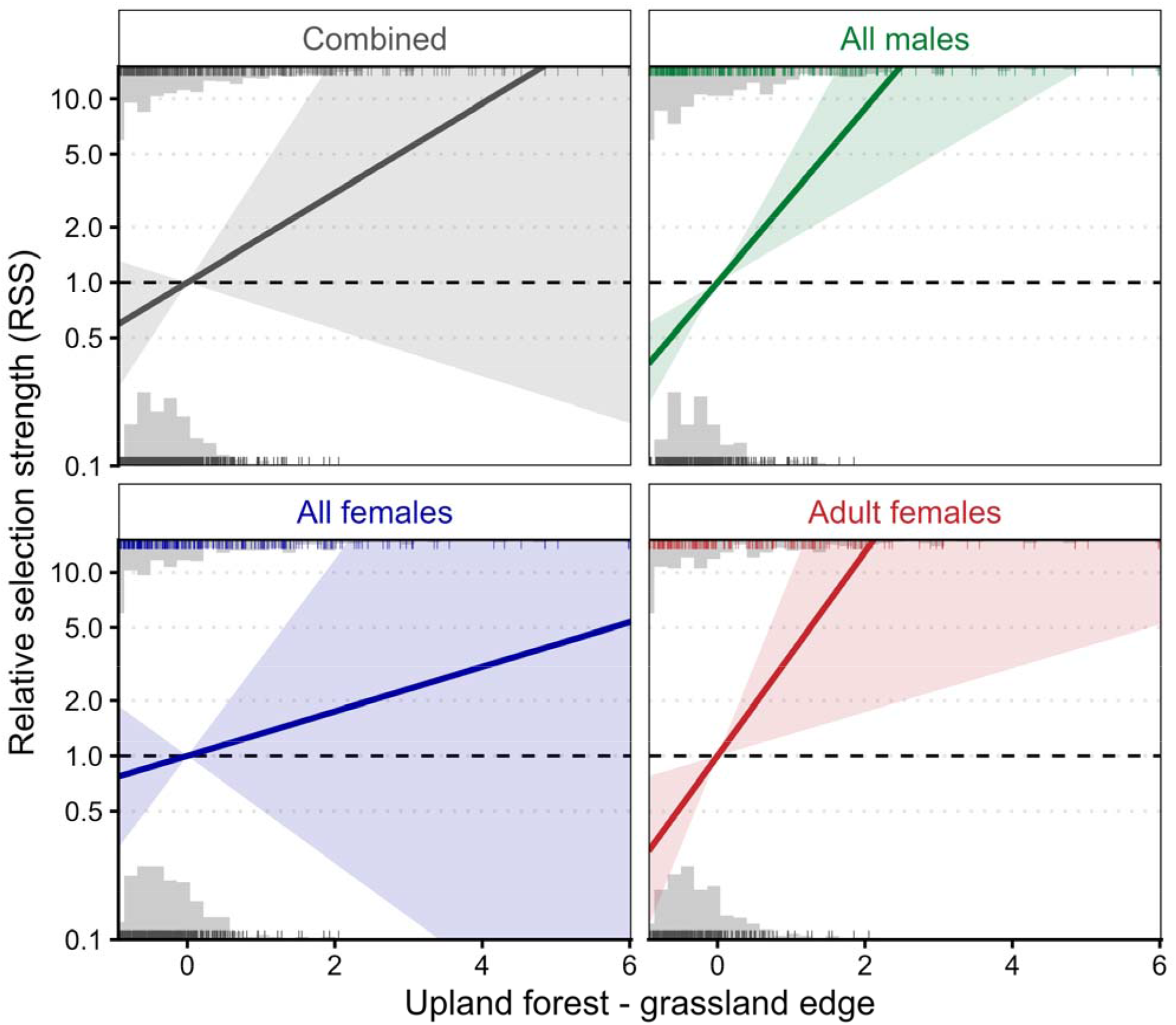
Resource selection response curves for upland forest–grassland edge across four bobcat demographic cohorts. The y-axis displays Relative Selection Strength (RSS) on a logarithmic scale. RSS quantifies the relative probability of use compared to average landscape availability, with the dashed horizontal line at RSS = 1 representing non-selection, and values > 1 indicating positive selection and values < 1 indicating avoidance. Solid lines represent the predicted model response for each cohort with shaded 95% confidence intervals. The x-axis represents the edge density metric centred and scaled to standard deviations with a mean of zero. Marginal histograms ticks mark the density and exact placement of used locations versus available background locations.

**Fig. 4.**
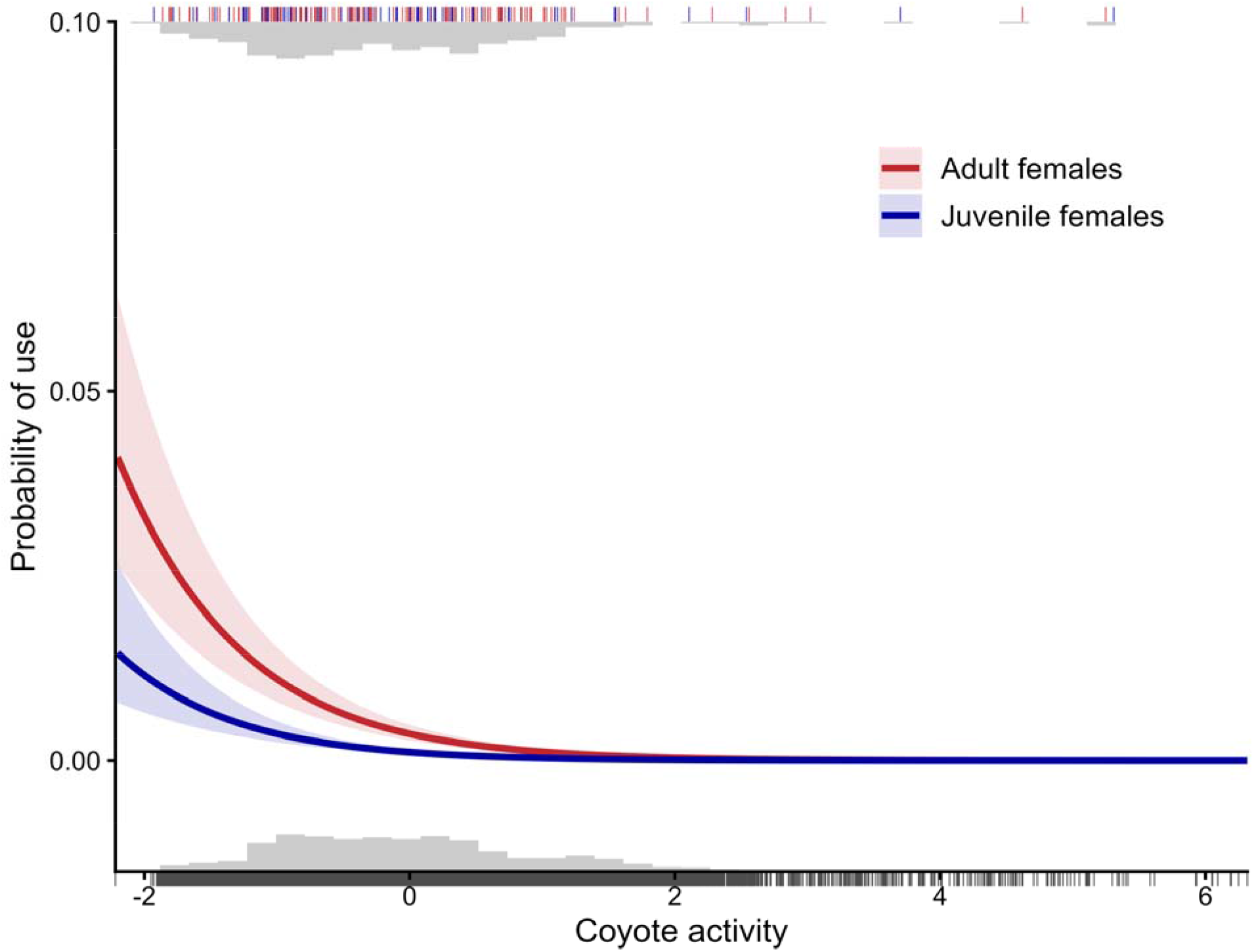
Adult and juvenile female bobcat response curves and 95% confidence intervals to coyote activity. The x-axis represents standardised coyote activity, with the mean set to zero. Both adult and juvenile females exhibited a negative response, indicating spatial avoidance of areas with high coyote activity. Marginal histograms and ticks illustrate the relative density and specific values of bobcat used versus available locations.

## Discussion

With few exceptions, our results indicate bobcat spatial ecology is tied to highly interspersed habitat mosaics motivated by their avoidance of expansive open habitats and the juxtaposition of woody vegetation with open canopy ecotones, consistent with our first hypothesis. Bobcat avoidance of high patch densities of open-canopy habitats confirms that even though those densities confer with them a higher density of ecotones, there is a threshold at which the advantage of high ecotone densities cannot offset the negatives associated with the lack of cover in these patches.

### Base requirements

Although their physiognomy may vary, extensive patches of even semi-natural open space is avoided by all bobcats with the sole exception of juvenile females at the landscape scale. The types of open spaces avoided includes large patches of pasture, proportion of the landscape consisting of grassland, and high densities of grassland and rowcrop patches. Whereas bobcats exhibited a clear avoidance of open habitats, likely due to a lack of cover, they strongly selected for woody vegetation – open ecotones (e.g. upland forest – grassland, woody wetland – rowcrop edges). This suggests that the value of open habitats to bobcats is conditional on adjacent woody structure. Counter-intuitively this avoidance of open areas may also explain the avoidance of mixed forest mean patch size by females. In Minnesota much of the mixed forest is characterized by a dominant conifer canopy with a sparse understory and limited prey (Buehler and Keith 1982). This may mirror the structural openness of a grassland at the ground level from the bobcat”s perspective, exposing them to risk while remaining prey poor.

### Centrality of ecotones

As our results support and felid evolutionary adaptations attest, ecotones are central to the spatial ecology of bobcats. The positive selection for edges across most models indicates a consistent reliance on the dense, structurally complex vegetation located at the juxtaposition of woody vegetation and canopy openings that can support high prey availability, while also providing escape cover and the sight-lines and concealment necessary for efficient prey capture (Fig. 3).

This mirrors the results of previous work focused on other felid species, including cougar (Elbroch et al. 2013; Knopff et al. 2014), jaguar (Alegre et al. 2024), as well as bobcats in other portions of their range (Abouelezz et al. 2018; McNitt et al. 2020). Notably, our results differ in that they identify the component ecotone habitats most important to different bobcat demographics, highlighting that all edges are not equally suitable.

For adult female bobcats, the selection of forest-edge ecotones paired with a landscape-scale avoidance of coyotes, severe winter weather, and expansive open habitats highlights a spatial response to a convergence of top-down and bottom-up pressures. Expansive patches of pasture, grassland, and rowcrop possess limited to no hunting cover or prey accessibility. Simultaneously, the lack of escape cover leaves bobcats exposed to competitors that are better equipped to exploit open habitats. Beyond competitors, this aversion to open habitats may reflect a response to anthropogenic risk. In the highly modified agricultural landscapes of the Midwest, large tracts of pasture and rowcrop increase exposure. For a medium-sized carnivore, navigating these open spaces significantly elevates the probability of human-caused mortality, whether through incidental detection or because open habitats provide easier points of access. Consequently, the adult female”s reliance on ecotones may not only be related to foraging, but a response to risk where dense forest edge provides thermal and escape refuge.

### Sex-based differences in spatial ecology

Differential habitat use by males and females likely represents a fundamental divergence in life history strategies owing to differing reproductive investment. Reproduction is an energetically expensive endeavour (Vaughn et al. 2011, p. 416), and disparate reproductive strategies magnify sex-based differences in selective pressures. In polygynous mating systems, males do not contribute to gestation or lactation, and only rarely to rearing, thereby producing unequal sex-based investment in reproduction; to maximize fitness, male felids prioritise mating opportunities whereas females seek to ensure offspring are recruited. Furthermore, because infanticide is a widespread male reproductive strategy (Balme and Hunter 2013; Steyaert et al. 2014; Palombit 2015), females may intentionally avoid males by using marginal habitats while young are dependent or outside of the breeding season (Pusey and Packer 1994; Wielgus and Bunnell 1995; Dejeante et al. 2024). To assess this further, we conducted a post hoc analysis of female space use relative to the predictions of male use using the top averaged male models. Across all scales, female use was positively associated with predicted male use (*β* > 0, *p* < 0.001), suggesting that sexual segregation is less important than resource availability.

Intraguild predation can alter the behavior and suppress populations of subordinate competitors (Palomares and Caro 1999). Despite documented coyote predation on juvenile and adult female bobcats, evidence for population-level suppression remains equivocal (Dyck et al. 2022). Our population-level models obscured sex-based divergence in spatial strategies relative to coyote activity, providing a plausible explanation for this ambiguity. Whereas our population-level model revealed a generalised avoidance of coyotes, the sex-specific models revealed this response was driven by female bobcats, and was entirely absent from the top male models. In sexually dimorphic species, the larger sex of the subordinate competitor can approach or even exceed the body size of the dominant competitor, a not uncommon occurrence for male bobcats and female coyotes (Donovan et al. 2011; Hinton et al. 2019), and could explain why male bobcats did not avoid coyote activity. To clarify inconsistent results from examinations of interspecific competition among carnivores, future research should consider how sexual size dimorphism might mediate interspecific competition.

Sexual size dimorphism also explains the female avoidance of freezing temperatures (*β* = -0.61, *p* < 0.001) and the counter-intuitive male selection for these same conditions (*β* = 0.62, *p* < 0.001) at the landscape level, albeit to slightly different metrics. Because the larger body mass of adult males confers a lower surface-area-to-volume ratio, they experience lower thermoregulatory costs than smaller females and juveniles. Therefore, the apparent selection by males possibly represents a greater physiological tolerance for severe cold, allowing them to prioritise the maintenance of their much larger territories regardless of freezing temperatures in early winter. Alternatively, the greater movement may reflect males increasing territorial defense in preparation of the oncoming breeding season in January to March (Crowe 1975).

At the local level, male bobcats exhibited strong selection for forested edges of rowcrops (*β* = 0.658, SE = 0.086, *p* < 0.001) while also exhibiting an even greater avoidance of rowcrop patch density (*β* = -2.750, SE = 0.442, *p* < 0.001), suggesting that the cumulative effect of greater fragmentation due to agriculture is negative and cannot be entirely offset by the added value of forest edge. One probable explanation for males selecting these edges is that they may provide access to prey which utilise crop residue from agricultural fields after harvest (Paisley et al. 1995; Delisle et al. 2024; Gilbertson et al. 2025) or the remnants of hunting bait and gut piles produce congregations of bobcat prey species (Bowman et al. 2015; Candler et al. 2023). That females did not also exhibit similar selection for forested agricultural edges may be due to the added risk that these ecotones pose, or that their lesser movement rates make them less able to detect these seasonal resource pulses.

### Ontogenetic shift

Consistent with our hypothesis, we detected a distinct ontogenetic shift in space use between juvenile and adult female bobcats, with adult females prioritizing the prey accessibility associated with forested-open canopy mosaics dominated by edge habitat and juvenile females prioritizing risk avoidance by limiting spatial overlap with adult females to intraspecific aggression, and exhibiting pronounced selection for vegetative structure and sensitivity to freezing temperatures and coyote activity. At the age of approximately 18 months, juvenile bobcats are beginning dispersal or territory establishment (Hughes et al. 2019), and are largely independent of maternal support and protection, making them the most vulnerable of all age groups, and this is reflected in their spatial ecology.

Examination of adult and juvenile female space use uncovered additional support for the supposition that combining demographic groups into a single category obscures fundamental differences in spatial ecology, in this case between adult and juvenile females. Whereas the upland forest–grassland ecotone was universally selected for local space use across all demographics, juvenile females exhibited spatial divergence at broader scales. Adult females (and all males) used semi-natural grassland edges heavily, while juvenile females selected for ecotones bordering otherwise highly avoided agricultural matrices, e.g. woody wetland–rowcrop (*β* = 0.50, *p* < 0.001) and upland forest–pasture (*β* = 0.34, *p* < 0.001) edges. Because large patches of pasture and rowcrop likely represent high risk components of the landscape, the juvenile selection for these edges suggests they are being competitively excluded from the less risky ecotones by dominant conspecifics. Adult females likely dominate the optimal, lower-risk natural ecotones, forcing subordinate juveniles to navigate the riskier, agricultural edges to find adequate prey and refuge. It is also possible that the use of these anthropogenic edges is the result of naivete in that juvenile females have not yet learned of the risk these ecotones pose (Loveridge et al. 2017; Thurfjell et al. 2017). Competitive exclusion may also explain the complexity of the juvenile female use of mixed forest. Across all demographics, juvenile females exhibited the only positive association to any mixed forest variable with their selection for mixed forest patch density (*β* = 4.28, *p* = 0.005). Although their use of high density of patches of mixed forest implies they provide a baseline level of suitability, juvenile female avoidance of large patches of mixed forests (*β* = -4.0, *p* < 0.001) suggests they may instead be using mixed forest to safely circumnavigate intraspecific aggression as they subsist in marginal habitats.

Both adult and juvenile females avoided coyote activity and freezing temperatures at the landscape level, supporting our hypothesis that animals respond to persistent, predictable environmental constraints such as regional climate gradients and dominant competitor distributions during home range establishment. However, the absence of this avoidance at the local scale contradicted our expectation of fine-scale spatial avoidance of localised, less predictable threats. We suspect this discrepancy is likely an artifact of spatiotemporal resolution rather than a true ecological absence. The grain of our data, which relied on harvest locations and track surveys, is robust for detecting broad, second-order landscape suitability, but lacks the spatiotemporal precision to capture the dynamic, transient behavioral avoidance that occurs at the scale of a single habitat patch (Jensen et al. 2024). Consequently, the lack of a statistically significant local-scale association does not preclude the existence of fine-scale risk mitigation, but rather our inability to detect it.

The ontogenetic divergence in risk tolerance between adult and juvenile female bobcats is not an isolated phenomenon, and similar patterns have been detected across other carnivore systems (Bauer et al. 2015; Schooler et al. 2025). In southern Africa, for example, juvenile male lions are relegated to marginal, low-precipitation habitats with lower prey abundance due to competitive exclusion by adults (Bauer et al. 2015). Similarly, smaller female grizzly bears, where body size strongly correlates with age, select for less productive habitats compared to larger females in Alaska (Schooler et al. 2025). We identified a comparable pattern in our system. Although both adult and juvenile females responded to the shared constraints of winter weather and interspecific competition at the landscape scale, juvenile females exhibited a lower baseline probability of use and greater sensitivity to risk. The steeper negative coefficients observed for juveniles in response to freezing temperatures (*β* = -0.75, compared to *β* = -0.61 for adults) and a lower multivariate baseline to coyote activity (Fig. 4) emphasise this heightened vulnerability, suggesting that the physiological and competitive costs of occupying this landscape are not equal for both age classes.

This heightened vulnerability to natural top-down and bottom-up pressures likely also impacted the ontogenetic divergence we observed in fine-scale edge selection. Forced to prioritise structural refuge from severe weather and intraguild predators, all while subjected to competitive exclusion from semi-natural ecotones by dominant adults, juvenile bobcat options are limited. And as a result, juvenile females must use the structurally dense but anthropogenically risky fringes of agricultural matrices such as woody wetland–rowcrop and upland forest– pasture ecotones. Intriguingly, while female avoidance of these constraints was only evident at the landscape scale, our inability to detect a response at finer scales likely reflects fine-scale behavioral flexibility. At the home range and local scales, females may exploit thermal and competitive refugia to buffer against transient conditions. It is probable that coarse-scale avoidance only becomes necessary when the severity or persistence of these stressors exceeds a critical threshold, rendering local refugia insufficient to offset associated physiological and survival costs.

## Conclusions

Animal resource selection emerges from a spatial negotiation between risk and reward. Our findings demonstrate that this trade-off is not a uniform, species-level trait, but is instead a dynamic, scale-dependent strategy shaped by sex and ontogeny. While baseline selection for foraging availability, such as found in upland forest–grassland ecotones, was conserved across the population, both the identity of the primary environmental drivers and the direction and magnitude of the spatial response to top-down and bottom-up constraints were dictated largely by an individual”s demographic stage.

Ultimately, our results highlight that the assumption of a single, population-wide pattern of habitat selection obscures the heterogeneity of carnivore habitat selection (Conde et al. 2010; Elliot et al. 2014; Bauer et al. 2025; Schooler et al. 2025; Zarzo-Arias et al. 2025). If conservation and management strategies rely solely on population-level responses, they risk missing the specific spatial resources critical for survival and recruitment (Adams et al. 2017) for less resilient demographies. Therefore, landscape management for solitary carnivores must shift from broad simplifications of species habitat needs to bespoke prescriptions that accommodate the divergent requirements of demographic groups.

